# Catastrophes, connectivity, and Allee effects in the design of marine reserve networks

**DOI:** 10.1101/750448

**Authors:** Easton R. White, Marissa L. Baskett, Alan Hastings

## Abstract

Catastrophic events, like oil spills and hurricanes, occur in many marine systems. One potential role of marine reserves is buffering populations against disturbances, including the potential for disturbance-driven population collapses under Allee effects. This buffering capacity depends on reserves in a network providing rescue effects, with a potential trade-off in spacing between individual reserves to balance rescue via connectivity and independence of disturbances experienced. We use a set of population models to examine how dispersal ability and the disturbance regime interact to determine the optimal reserve spacing. We incorporate fishing in a spatially-explicit model to understand the consequences of objective choice (i.e. conservation versus fisheries yield) on the optimal reserve spacing. We show that the optimal spacing between reserves increases when accounting for catastrophes. This result is accentuated when Allee effects interact with catastrophes to increase the probability of extinction. We also show that classic tradeoffs between conservation and fishing objectives disappear in the presence of catastrophes. Specifically, we found that at intermediate levels of disturbance, it is optimal to spread out reserves in order to increase population persistence and to maximize spillover into non-reserve areas.

## 1 Introduction

Marine protected areas (MPAs), including no-take marine reserves (Lubchenco et al. 2003), are increasingly being used as a form of ecosystem-based management (Wood et al. 2008). Management goals of MPAs range from promoting sustainable fisheries to conserving biodiversity (Leslie 2005; Gaines et al. 2010). Potential outcomes of MPAs include increases in biomass and population size as well as spillover to harvested areas (Lester et al. 2009; Gaines et al. 2010; Baskett and Barnett 2015). Beyond individual reserves, networks of reserves can connect over larger areas given the long-distance passive dispersal of many marine organisms at early life history stages (Kinlan and Gaines 2003). Even in situations where each single reserve is not self-sustaining, network persistence can still be possible (Hastings and Botsford 2006). This overall network persistence depends on the specific size of reserves and the spacing between them (Botsford et al. 2001; 2003; Gerber et al. 2003; Gaines et al. 2010).

The outcome and optimal design of reserve networks can depend on environmental variability, including disturbances (Halpern et al. 2006; Cabral et al. 2017; Aalto et al. 2019). Marine systems are inherently variable due to seasonal forces affecting temperature and upwelling, disturbances, and longer term cycles including El Niño (Fiedler 2002; Doney et al. 2012; White and Hastings 2019). Several of these factors (e.g. storms, marine heat waves) are expected to become stronger or more variable in the future because of climate change (Bender et al. 2010; Oliver et al. 2018). This variability inevitably affects the population dynamics and distributions of many organisms. In addition, environmental variability or uncertainty can alter the optimal management strategies (Halpern et al. 2006). For example, temporal variability can increase the role of marine reserves in population persistence (Mangel 2000). Further, the optimal reserve selection depends on the scale of stochasticity, either local demographic noise or regional disturbances, in the system (Quinn and Hastings 1987).

As an extreme form of environmental variability, disturbances (i.e. rare events or catastrophes) in marine systems can include marine heat waves, hurricanes, oil spills, hypoxia, economic shocks, and disease outbreaks (Mangel and Tier 1994; Allison et al. 2003; Game et al. 2008b; Aalto et al. 2019; Cottrell et al. 2019; White et al. 2020). Given disturbances, the individual reserves within a network can act as an insurance policy for other reserves, i.e. a portfolio effect across reserves (Wagner et al. 2007; Pitchford et al. 2007). The idea of a spatial insurance policy is one of the many arguments raised for allocating protection in more than one location in the “single large or several small” (SLOSS) debate that originated in the terrestrial reserve literature (Fahrig 2013). For instance, Kallimanis et al. (2005) used a spatially-explicit metapopulation model to show that, while a single large area was favored with random disturbances, when disturbances were spatially autocorrelated this relationship no longer held. Similarly, with a more tactical model, Helmstedt et al. (2014) showed that several, smaller predator exclusion patches were favored in the presence of catastrophes. However, findings from the terrestrial realm may not translate to the marine setting given the greater potential for passive dispersal, greater permeability of reserve boundaries, and fishing rather than habitat loss as the primary anthropogenic impact (Carr et al. 2003; Halpern and Warner 2003).

For marine reserve networks, disturbances can increase the amount of protection needed to achieve management goals (Allison et al. 2003) and increase the optimal distance between reserves (Wagner et al. 2007). Reserves placed too close together will potentially be affected by the same disturbance event simultaneously, reducing the opportunity for one reserve to “rescue” another (sensu Brown and Kodric-Brown 1977). Conversely, reserves placed too far apart also reduce the probability of successful dispersal between protected areas. However, with certain forms of disturbances, such as disease, it might be better to have reduced dispersal between areas (Sokolow et al. 2009; Kough et al. 2015). Thus, when incorporating catastrophes into reserve network design, spatial scales of species movement and disturbance become important (Quinn and Hastings 1987; Wagner et al. 2007; Blowes and Connolly 2012) and can alter the optimal size and spacing of reserves within a network.

Disturbances can have particularly strong effects in systems that exhibit alternative stable states (Paine et al. 1998; Fabina et al. 2015; Dennis et al. 2016) by shifting a system from one state to another (Scheffer et al. 2001; Scheffer and Carpenter 2003). When a system moves from one state to another, hysteresis impede recovery of the original state. One source of alternative stable states within populations are strong Allee effects, which arise from positive density-dependence at low densities (Dennis et al. 2016). Allee effects have been well-documented for a number of species, including marine fishes (Hutchings 2013), and increase extinction risk (Dennis et al. 2016). In systems with alternative stable states, including Allee effects, MPAs can, theoretically, decrease the likelihood of state shifts compared to traditional fisheries management by protecting a population or community further from thresholds within their boundaries that then, through dispersal, can be a source for rescue effects in harvested areas (Barnett and Baskett 2015; Aalto et al. 2019).

We build a series of models to quantify the optimal distances between reserves affected by disturbances. We first consider a set of simple population models of two-patch systems. This is analogous to Wagner et al. (2007), but with the added realism of density-dependent recruitment which can be an important buffer against environmental variability (Botsford et al. 2015). We then extend this framework to investigate how Allee effects and dispersal ability interact with the disturbance regime characteristics (e.g. frequency and magnitude) to affect the optimal spacing between reserves. This interaction occurs as disturbances have the potential to push populations below an Allee threshold, causing a shift to an unfavorable alternative state. This builds on work by Aalto et al. (2019), who demonstrate the buffering capacity of MPAs given disturbance and Allee effects, to explore the question of optimal spacing in a reserve network. Finally, we examine a spatially-explicit, multiple patch model of a coastline to explore the robustness of conclusions from the two-patch scenario and the role of fishing on network design. Including outside-MPA harvest also allows exploration of the optimal spacing of MPAs for both fisheries and conservation goals.

## 2 Methods and models

### 2.1 Two-patch model

First, we use a two-patch, coupled Beverton-Holt model. In patch *i, N*_*i*_(*t*) is the population size at time *t*. During each time step, we model three sequential events: 1) production, 2) dispersal, and 3) disturbance (Fig. 1). We incorporate density-dependence in production with a Beverton-Holt function, *G*_*i*_, where *r*_*i*_(*t*) is the growth factor and *K*_*i*_ is the carrying capacity, i.e., maximum value where population growth saturates. We describe *r*_*i*_ as a normal distribution to allow temporal variability in the growth factor with mean *µ*_*r*_ and variance 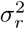. We assume both patches are reserves where no fishing occurs. This implicitly contains a “scorched earth” assumption that fishing removes all biomass outside of these two patches (Botsford et al. 2001). In addition to negative density dependence arising from carrying capacity *K*_*i*_, we incorporate the potential for Allee effects with the factor *ω* where an Allee effect is present (initially concave relationship between *G*_*i*_ and *N*_*i*_(*t*)) for *ω* > 1 (Fig. 1b). The growth function for each patch is then:

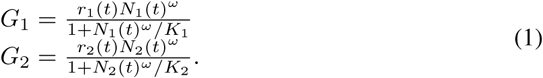

**Figure 1:**
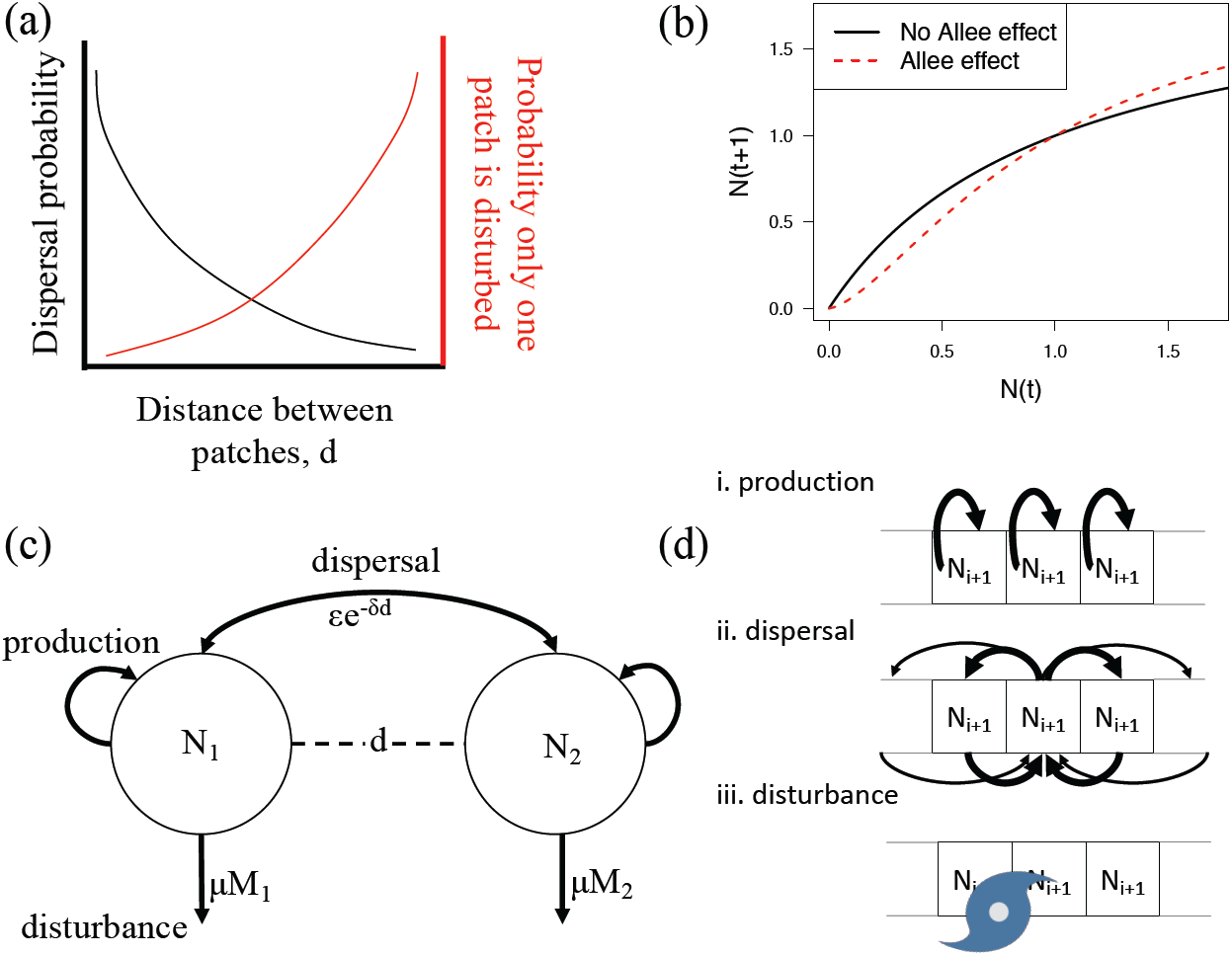
(a) Conceptual trade-off between successful dispersal probability and probability that a given disturbance only affects a single patch locally for different distances between patches, *d*. (b) Production functions for population size from year t to year t+1 without (*ω* = 1) and with (*ω* = 1.5) an Allee effect. (c) Model diagram for the two-patch reserve scenario. Here *N*_*i*_ is the abundance in patch *i, µM*_1_ is the mortality caused by disturbance, and *ϵe*^−*δd*^ is the probability of successful dispersal. The arrows pointing out and back towards the same patch denote the recruitment process. (d) Model for *n*-patches cycles through production within patches (given by the self-arrow), dispersal between patches (denoted by arrows connecting patch), and a disturbance centered on the middle patch that also affects the nearby patch to the left.

We connect the two patches via the dispersal, with fraction *ϵ* of dispersing individuals (Fig. 1c). Similar to a Laplacian (double-exponential) dispersal kernel, we model dispersal success as an exponentially decaying function of distance, *d*, between reserves at a rate *δ* to yield the post-dispersal population sizes:

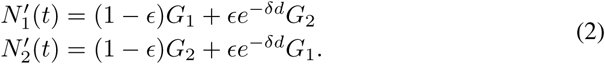

Finally, we model disturbance in each patch, *M*_*i*_, as a binomial distribution with probability, *p*_*i*_. If disturbance affects one patch, the other patch has a probability *e*^−*γd*^ of also being affected by the disturbance, where *γ* is a shape parameter. Therefore, the disturbance occurrences are:

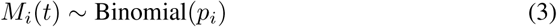

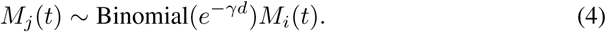

A disturbance causes density-independent mortality, *µ*, for the entire patch, This mortality and 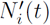 give the number of individuals the following year is:

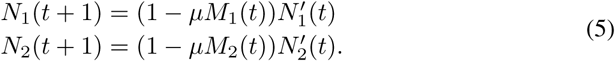

This then presents a trade-off in reducing patch distance to increase colonization potential, but increasing patch distance to reduce the probability of disturbance in both patches simultaneously (Fig. 1a).

### 2.2 *n*-patch model

We extend our two-patch example to a model with *n* patches (Fig. 1d). We focus on a coastline system with a simple one-dimensional landscape (Fig. 1d). Thus, we examine a set of discrete, contiguous patches with some patches designated as reserves (i.e. the reserve fraction of the coastline) and other patches exposed to fishing (Botsford et al. 2001). The within-patch production function is the same as the two-patch model, but with added fishing mortality in unprotected areas after dispersal occurs. In this spatially explicit model, dispersal occurs according to a geometric distribution, and we scale disturbance by an exponential distribution around a focal patch. We assume that any individuals that disperse outside the *n* patches do not survive. We describe this *n*-patch fully in the supplementary material.

### 2.3 Reserve network optimization

For the two-patch model, we examined all possible distances (at integers up to 100 units apart) between patches. With the default dispersal and disturbance parameters, distances of greater than 100 units were never optimal. The optimal spacing was the distance that maximized population persistence (proportion of runs where the ending total population size is positive). Stochasticity was low in the default model (Table 1), so only 100 runs of each parameter combination were needed.

**Table 1.** Parameter notation, description, and default values for the two-patch model. As a sensitivity analysis, several parameters (e.g. *d, ω*) are varied in the Figs. 2,3,S1-S3. The *n*-patch model is described more fully in the supplementary material.

| Notation | Description | Default value (units) |
| --- | --- | --- |
| $r_i(t)$ | growth factor of patch $i$ at time $t$ described as a normal distribution | |
| $\mu_r$ | mean of growth factor normal distribution | 2 |
| $\sigma_r^2$ | variance of growth factor normal distribution | 0.25 |
| $K_i$ | carrying capacity for patch $i$ | 1 |
| $\omega$ | Allee effect parameter | 1 for no Allee effect or $>1$ for Allee effect |
| $\epsilon$ | fraction of dispersing individuals | 0.5 |
| $d$ | distance between patches | 0-100 (spatial units) |
| $\delta$ | dispersal success parameter (larger $\delta$ indicates lower survival) | 0.01 (1/spatial unit) |
| $p_i$ | probability of disturbance | 0.02 |
| $\gamma$ | decay parameter for probability that disturbance in one patch affects the other | 0.1 (1/spatial unit) |
| $\mu$ | density-independent mortality from disturbance | 0.8 |

In the *n*-patch model, we examined two different management goals for determining the optimal reserve configuration: (1) maximize the probability of population persistence, measured as the proportion of runs that end in positive population size and (2) maximize the total fisheries yield, which is the total catch over the entire time series. We then examine different reserve configurations to determine the optimal spacing between reserves. For example, a system with 12 discrete patches, including three reserves, could be NNN**R**N**R**NN**R**NNN where R and N represent reserve and non-reserve sites, respectively. In this example, the average spacing between reserves is 2.5. We examine all the possible orderings of reserves and non-reserves. Thus, we systematically examine all possible reserve networks. This is possible when the number of patches is small. We limit the fraction of area that can be placed in reserves, specifically 20% of the patches (we test the robustness of our results to different parameter values in Fig. S4). For each network configuration, we run 100 simulations to account for stochasticity in model runs generated by the random disturbance locations and frequencies (example dynamics in Fig. 2). We then calculate the mean value of persistence and total fisheries yield across these simulations to determine the performance of that reserve network. We chose a set of parameters that are biologically-realistic compared to other fisheries (Britten et al. 2016) and lead to a space where certain key dynamics (e.g. trade-off between catastrophes and connectivity) are relevant. We examine the sensitivity of our results to changes in each of the parameters (Figs. S1-S3).

**Figure 2:**
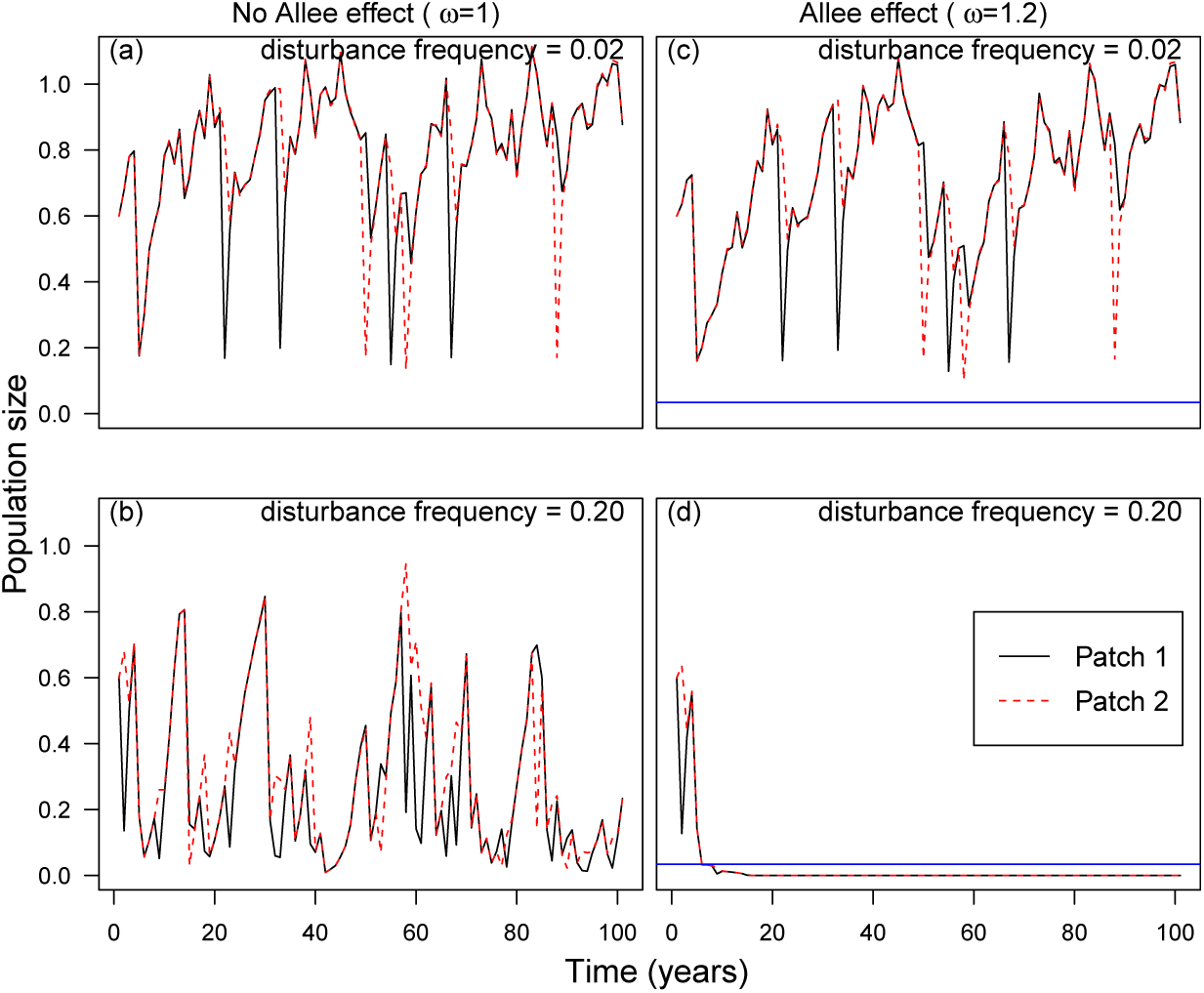
Example two-patch model simulation, each line denotes a different patch. The four panels represent different combinations of disturbance frequency and the presence (*ω* > 1) or absence of an Allee effect. The horizontal line denotes the Allee threshold. The simulation is with the following model parameters: *r* = 2.0, *K* = 1, *δ* = 0.01, *ϵ* = 0.5, and distance between patches of 40.

## 3 Results

We find that catastrophes can increase the optimal spacing between a pair of reserves (Fig. 3a). To maximize population persistence, some spacing between patches is optimal (Fig. 3a) whenever catastrophes are present. However, as the disturbance frequency or magnitude increases, the population size (and population persistence) decreases. Thus, the optimal spacing between reserves increases with disturbance frequency, but when disturbances are too frequent the population goes extinct from the catastrophes (Fig. 3a). Without accounting for catastrophes (when disturbance frequency is zero), populations can only go extinct because of temporal variability in the growth rate. In this case, the optimal strategy to maximize population persistence is to place the reserves adjacent to each other (Fig. 3a).

**Figure 3:**
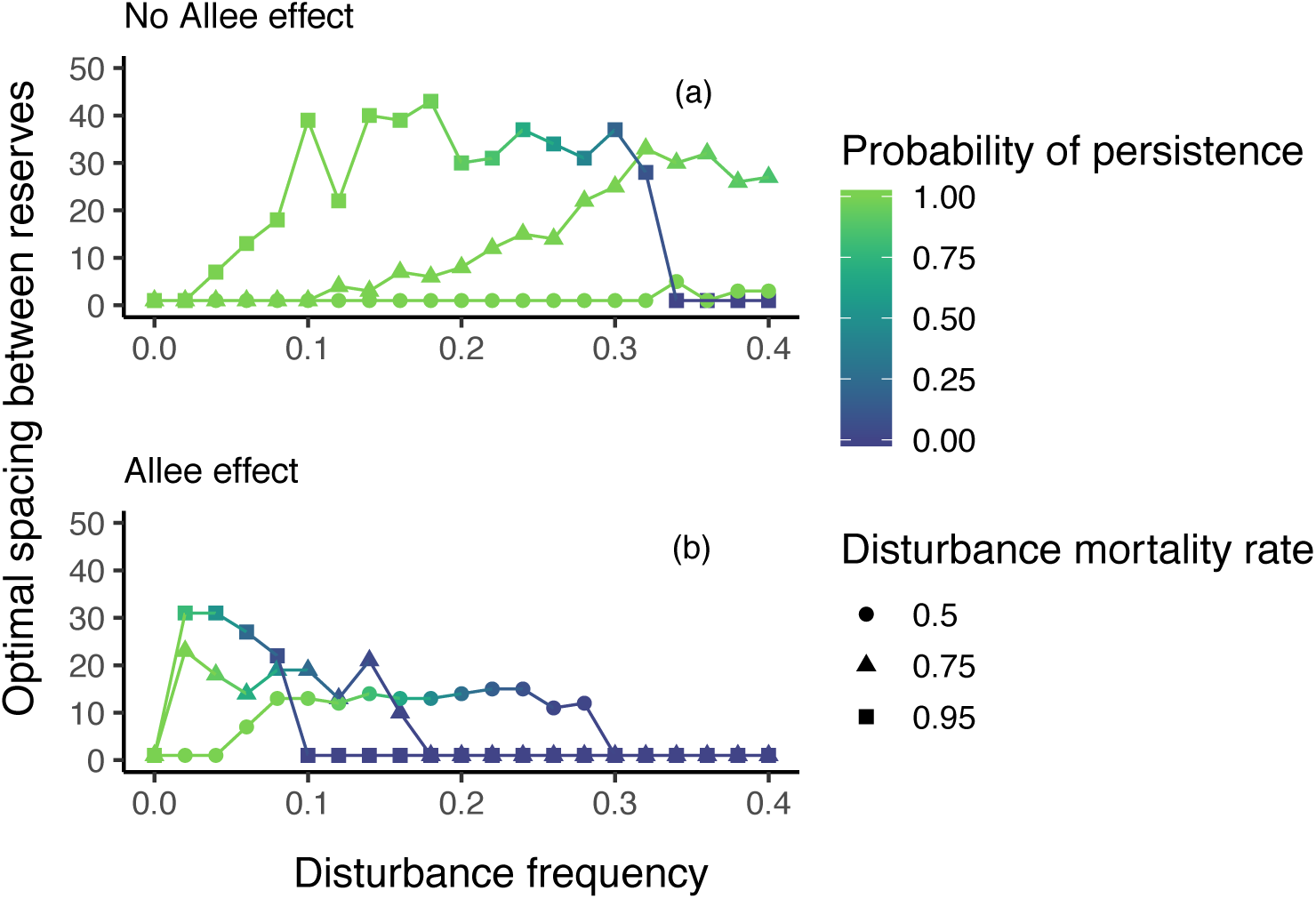
Optimal spacing for different disturbance frequencies and mortality rates. Optimal spacing for the Beverton-Holt model (a) without (*ω* = 1) and (b) with an Allee effect (*ω* = 1.2). Each line indicates a different disturbance mortality rate. The color of each point is the fraction of runs where the end population size was non-zero, a measure of persistence. These simulations are with the same model parameters as in Fig. 2.

The presence of an Allee effect interacts with catastrophes to increase the risk of extinction (Fig. 2c,d), which decreases optimal spacing between reserves (Fig. 3b). In addition, disturbance level, once present, and mortality rate have minimal effect on optimal spacing when we account for Allee effects (Fig. 3b).

Intuitively, the optimal distance between reserves increases with longer-distance dispersal (Figs. 4). Dispersal has the strongest effect on reserve spacing at low disturbance probabilities and consequently small inter-patches distances are optimal unless there is longer-distance dispersal (Fig. 4). Conversely, when disturbances are likely to occur in both patches simultaneously it is optimal to space reserves closer together (Fig. 4). This happens because disturbances are large enough to drive the entire reserve network to extinction regardless of the organism’s dispersal ability.

**Figure 4:**
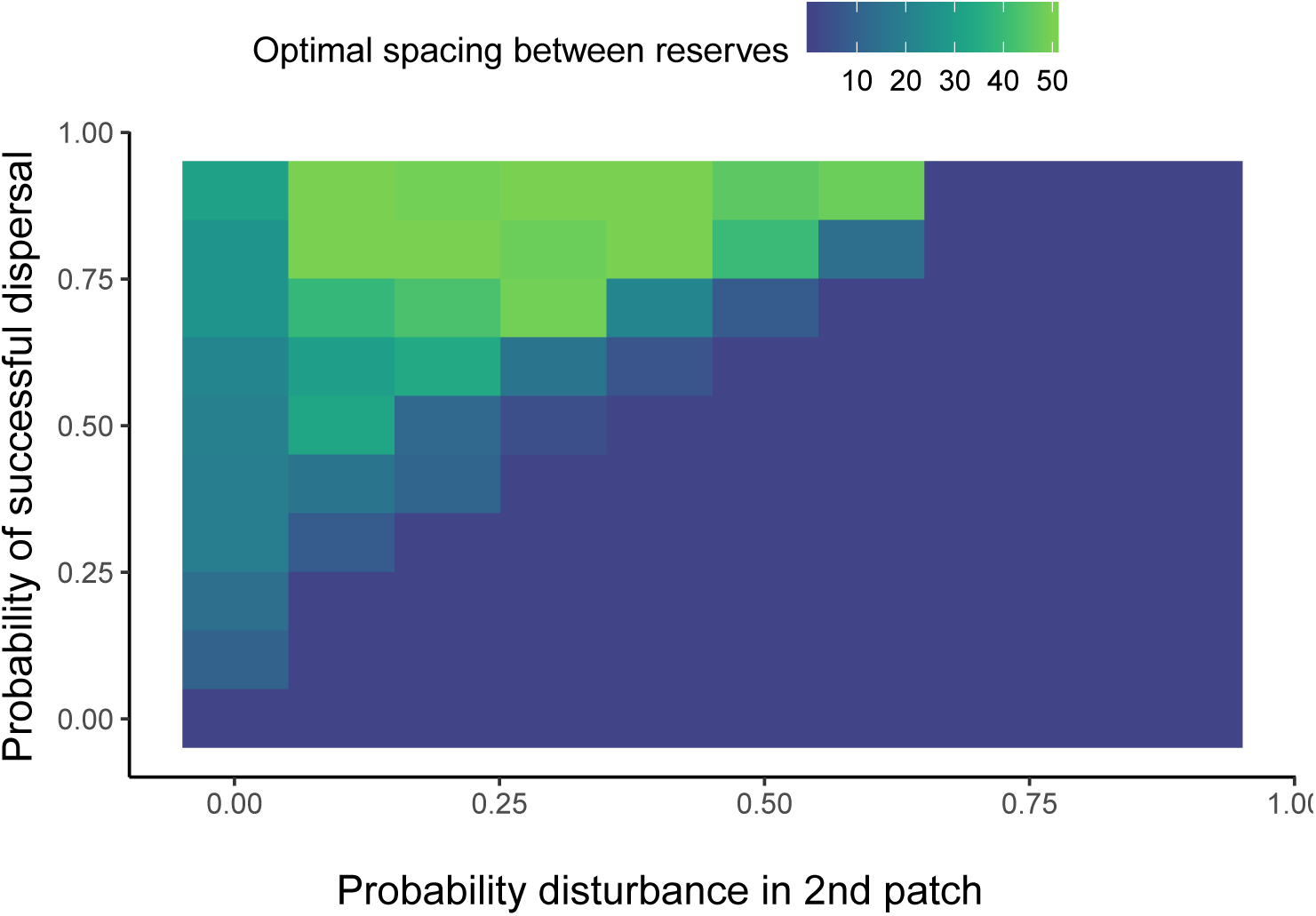
The optimal spacing between reserves for population persistence given different combinations of successful dispersal probability (to a patch 50 units away) and the probability that a disturbance in one patch affects the nearby second patch (at a distance of 50 units away). Simulations here are for a two-patch model with an Allee effect (*ω* = 1.2) and the same parameter values as in figure 2.

### 3.1 *n*-patch simulation model

Optimal spacing for fisheries and conservation goals diverge under no disturbance and converge with disturbance (Fig. 5). Without disturbances, maximizing fisheries spillover can be achieved through spreading out reserves. However, in order to maximize population persistence, it is optimal to have no reserve spacing, i.e. a single, larger reserve. For moderate disturbance frequencies, the optimal spacing between reserves can be nearly the same for the fishing and persistence objectives (Fig. 5). With increasing fraction of coastline protected, disturbances are less important and reserves can be placed closer together when maximizing population persistence (Fig. S4).

**Figure 5:**
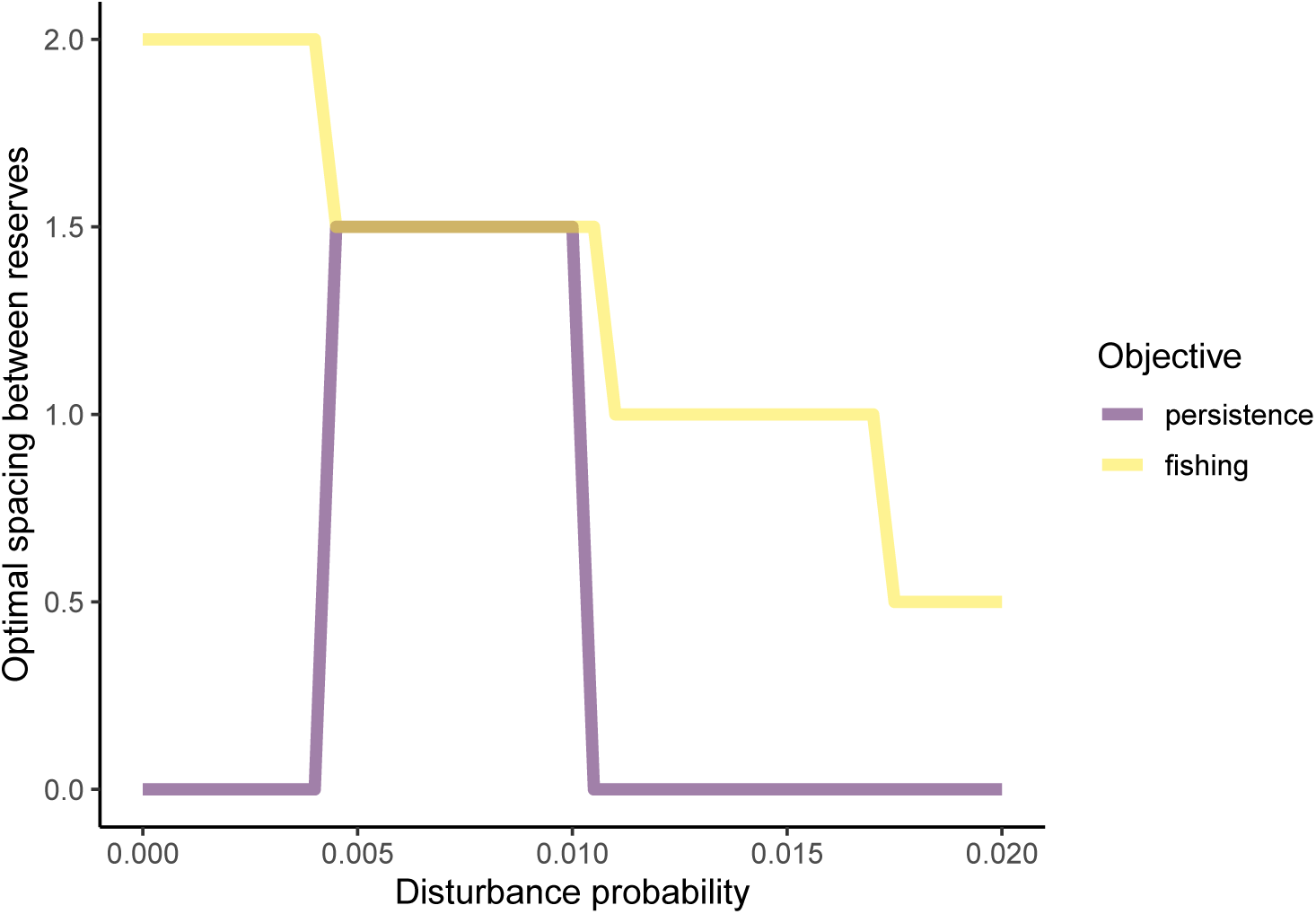
Optimal spacing between reserves (in an *n*-patch model) versus the probability of disturbance for two different objective functions. The parameter values used are: *δ* = 0.7, *ω* = 1.2, r = 3, and *µ* = 0.95.

All of our above results include a “scorched earth” assumption, where there is 100% mortality for individuals outside of the reserves. When we relaxed this assumption, we found that, for reduced fishing pressure, persistence was maximized when reserves were placed close together at low disturbance frequencies (Fig. 6). At high disturbance frequencies, this relationship reverses and reduced fishing pressure increase the optimal spacing between reserves (Fig. 6). This happens because fishing pressure outside reserves reduces the capacity for the system to recover from catastrophes.

**Figure 6:**
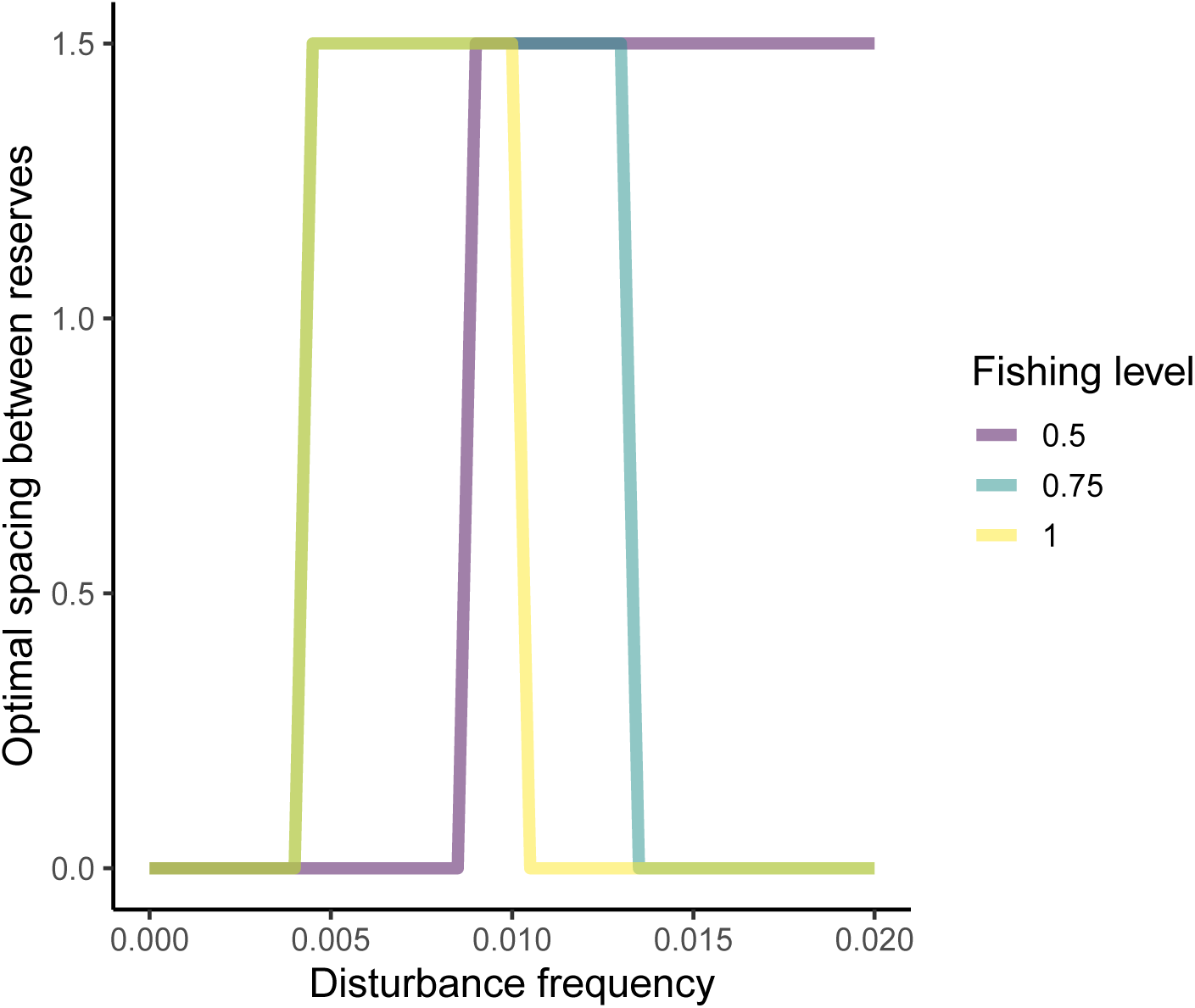
Optimal spacing between reserves (in an *n*-patch model) to maximize persistence for different disturbance frequencies and fishing levels outside of reserves (fraction of population harvested in each patch). The parameter values used here are: *δ* = 0.7, *ω* = 1.2, r = 3, and *µ* = 0.9.

## 4 Discussion

Disturbances can increase the spacing in reserve networks to achieve an objective of maximizing persistence (Figs. 3,5). The optimal distance is a trade-off between the disturbance size and frequency as well as the dispersal ability of the organism. These results are in line with that of Wagner et al. (2007), who found that persistence was maximized at intermediate inter-reserve distances because of the same set of tradeoffs. We expand their findings with our consideration of density dependence and fisheries outside reserves, where we find that persistence is not always maximized with spacing between reserves. Instead, the exact spacing depends on the specific disturbance regime and the life history (i.e. dispersal) of the species. Our results are most analogous to situations where disturbances directly cause additional mortality, for instance hurricanes and oil spills (Allison et al. 2003). If disturbances are not frequent, or only have a marginal effect on survival rates, it will still be optimal to place reserves nearby one another to maximize connectivity (Figs. 3,5).

The benefit to spacing out reserves, for potential post-disturbance rescue across a reserve network, is most relevant to long-distance dispersers, and short-distance dispersers do not experience this benefit in our model (Fig. 4). In the extreme, our results with short-distance dispersal approach the model without dispersal by Game et al. (2008a), who found that protecting larger reserves was more optimal than smaller, spread out reserves when catastrophes are present. McGilliard et al. (2011) explicitly modeled dispersal, making their models most akin to our own. In the presence of disturbances, they found that persistence was unlikely if only a marine reserve was the only spatial management strategy. We build on this work by showing that persistence is possible using a network of reserves, as opposed to a single large reserve. Thus, we see that if the probability of successful dispersal is higher than the probability of disturbance affecting reserves simultaneously, then it is always optimal to spread out reserves when disturbances are present (Fig. 4).

Including Allee effects in our model, decreased persistence and increased the optimal spacing between reserves (Figs. 2d,3b). A strong Allee effect creates alternative stable states with a tipping point at the Allee threshold. Catastrophes interact with Allee effects by causing a regime shift to an alternative stable state (Dennis et al. 2016). Thus, it is necessary to spread out reserves farther in cases where Allee effects are present (Fig. 3b). This increases the likelihood that acting as an insurance policy for one another plays a role in persistence.Our findings are in line with past theoretical findings on catastrophes and the distance between patches within a metapopulation (Blowes and Connolly 2012; Dennis et al. 2016) showing the robustness of our results. Thus, designing reserve networks with catastrophes in mind will be particularly relevant for species that experience mate limitations at low density or that rely on larger populations for survival (Courchamp et al. 1999). Specific examples include species like Atlantic cod (*Gadus morhua*) or Alaskan walleye pollock (*Theragra chalcogramma*) where Allee effects have been identified (Hutchings 2013). Aalto et al. (2019) studied a population of green abalone (*Haliotis fulgens*) in Baja California Sur, Mexico which experiences an Allee effect through recruitment failure at low population densities. They show that a reserve network can increase persistence for abalone when they are exposed to catastrophic events. Our work builds on this by focusing on the specific design of reserve networks.

We found that catastrophes can lead to alignment in the optimal design of reserves networks for objectives that would otherwise trade off with each other. In analyzing different reserve objectives, we found that, at intermediate disturbance levels, the optimal spacing between reserves was the same for both maximizing fishing and maximizing persistence (Fig. 5). In comparison, we find that without catastrophes (disturbance frequency is zero), large, single reserves maximize persistence and smaller, spread out reserves maximize fishing as with other papers without disturbance (Gaines et al. 2010). This conflict arises because spreading out reserves maximizes spillover from reserves to fished areas, but reduces within-reserve protection. However, when accounting for catastrophes in our models, disturbances lead to an optimal reserve design where reserves are spread out in order to decrease the probability of synchronous collapse of all reserve areas and therefore increase the probability of persistent spillover.

Considering the management outside of reserves, we found that decreasing fishing pressure outside reserves can decrease the optimal spacing between reserves (Fig. 6). This happens because of higher abundance outside reserves, reducing their buffering effect. In line with our results, McGilliard et al. (2011) found that fishing management outside a reserve was important to maximize persistence. In their work, this occurred because reduced fishing pressure allowed for higher abundances to be maintained, allowing for a spatially diffuse population. The spatially diffuse population was then less susceptible to local disturbances. Importantly, they only study the effect of the presence or absence of a single reserve. Our results build on this work by showing how the optimal reserve network design interacts with fishing pressure.

As with any model, we make a number of simplifications. First, the patch connectivity would have to be determined using an approach like a regional oceanic modeling system (ROMS) model or other connectivity data (Shchepetkin and McWilliams 2005; Watson et al. 2012). A more realistic connectivity model might alter the optimal spacing between reserves given specific sink-source dynamics (Crowder et al. 2000) and variability in connectivity between years (Williams and Hastings 2013). More specifically, the relationship between the spatial scale of dispersal and disturbance interact to determine optimal reserve configurations (Quinn and Hastings 1987; Kallimanis et al. 2005); this might be especially true in sytstems with landscapes more complex than a simple linear coastline (Moloney and Levin 1996). Further, our model includes local dispersal and local catastrophes. Actual connectivity data could decouple these processes if dispersal is on much larger scales than catastrophes. We ignore age structure, which can have a buffering effect to disturbances if age classes are affected deferentially. In a system with sedentary adults, including age structure might increase the production within reserves. This, in turn, concentrates the population, increasing the importance of spacing reserves in the presence of catastrophes. Furthermore, disturbance likelihood can be spatially and temporally heterogeneous, where some parts of the coastline might experience lower-frequency disturbance (Game et al. 2008a; Mellin et al. 2016). Game et al. (2008a) found that for realistic rates of cyclones affecting the Great Barrier Reef, persistence was maximized by protected subsets of the reef, the specific subsets depending on the cyclone frequency and the post-disturbance recover rate. Thus, for our results on optimal spacing, heterogeneous disturbances could cause variable spacing between reserves or prioritizing of certain regions. Although there exists high quality data on the location and frequency of different disturbances (e.g. US National Hurricane Center https://www.nhc.noaa.gov/data/; Coral Reef Watch https://coralreefwatch.noaa.gov/), connecting these data to population dynamics also requires knowledge about the demographic effects (e.g. mortality rates of corals for different cyclone severity) these have on marine organisms. Further, we only focused on disturbances that act independent of the population dynamics (e.g. hurricanes). Disturbances, such as diseases, that can negatively affect population dynamics through connectivity (e.g., disease spread from dispersal) then increase the spacing expected to optimize population persistence (Sokolow et al. 2009; Kough et al. 2015). Lastly, our study focused on a single species. For multi-species systems, species can have different life histories, including their dispersal abilities and susceptibility to disturbances. This could lead to different reserve networks for multi-species systems (D’Aloia et al. 2017), including scenarios where variable spacing or sizes of reserves might be optimal. Thus, future work should examine system-specific life-history traits and disturbance regimes as well as multiple species.

## 5 Author Contributions

ERW, MLB, and AH conceived the project idea and analyses; ERW designed and wrote the models. All authors contributed critically to the drafts and gave final approval for publication.

## 6 Acknowledgments

ERW was partially supported by a National Science Foundation Graduate Fellowship. AH was supported by grant DMS-1817124. We would like to thank members of the Ecological Theory group at the University of California, Davis for their insight.

## 7 Supporting Information

In the supporting material, we provide a description of the *n*-patch model and additional figures. All code and data are available at https://github.com/eastonwhite/MPA-disturbances.

## Notes

### Competing Interest Statement

The authors have declared no competing interest.

### Summary of Updates

This manuscript has been revised with new figures and supplementary material.

## References

E. A. Aalto, F. Micheli, C. A. Boch, J. A. E. Montes, C. B. Woodson, and G. A. D. Leo. Catastrohic mortality, allee effects, and marine protected areas. The American Naturalist, 193(3):391–408, 2019. doi: 10.1086/701781.

G. W. Allison, S. D. Gaines, J. Lubchenco, and H. P. Possingham. Ensuring Persistence of Marine Reserves-Catastrophes Require Adopting an Insurance Factor. Ecological Applications, 13(1):S8–S24, 2003.

A. Barnett and M. L. Baskett. Marine reserves can enhance ecological resilience. Ecology Letters, 18(12):1301–1310, 2015. doi: 10.1111/ele.12524.

L. Baskett and L. A. Barnett. The Ecological and Evolutionary Consequences of Marine Reserves. Annual Review of Ecology, Evolution, and Systematics, 46(1):49–73, 2015. ISSN 1543-592X 1545-2069. doi: 10.1146/annurev-ecolsys-112414-054424.

M. A. Bender, T. R. Knutson, R. E. Tuleya, J. J. Sirutis, G. A. Vecchi, S. T. Garner, and I. M. Held. Modeled Impact of Anthropogenic Warming on the Frequency of Intense Atlantic Hurricanes. Science, 327(5964):454–458, Jan. 2010. ISSN 0036-8075, 1095–9203. doi: 10.1126/science.1180568.

S. A. Blowes and S. R. Connolly. Risk spreading, connectivity, and optimal reserve spacing. Ecological Applications, 22(1):311–321, 2012.

L. W. Botsford, A. Hastings, and S. D. Gaines. Dependence of sustainability on the configuration of marine reserves and larval dispersal distance. Ecology Letters, 4(2): 144–150, 2001. ISSN 1461-023X. doi: 10.1046/j.1461-0248.2001.00208.x.

L. W. Botsford, F. Micheli, and A. Hastings. Principles for the design of marine reserves. Ecological Applications, 13(1):S25–S31, 2003.

L. W. Botsford, M. D. Holland, J. C. Field, and A. Hastings. Cohort resonance: A significant component of fluctuations in recruitment, egg production, and catch of fished populations. Marine science, no hay:1–7, 2015. ISSN 1206616369. doi: 10.1093/icesjms/fst176.

G. L. Britten, M. Dowd, and B. Worm. Changing recruitment capacity in global fish stocks. Proceedings of the National Academy of Sciences, 113(1):134–139, Jan. 2016. ISSN 0027-8424, 1091–6490. doi: 10.1073/pnas.1504709112.

J. H. Brown and A. Kodric-Brown. Turnover Rates in Insular Biogeography: Effect of Immigration on Extinction. Ecology, 58(2):445–449, Mar. 1977. ISSN 00129658. doi: 10.2307/1935620.

R. B. Cabral, B. S. Halpern, C. Costello, and S. D. Gaines. Unexpected management choices when accounting for uncertainty in ecosystem service tradeoff analyses. Conservation Letters, 10(4):421–429, 2017. ISSN 1044071060. doi: 10.1111/conl.12303.

M. H. Carr, J. E. Neigel, J. A. Estes, S. Andelman, R. R. Warner, and J. L. Largier. Comparing marine and terrestrial ecosystems: Implications for the design of coastal marine reserves. Ecological Applications, 13(sp1):90–107, Feb. 2003. ISSN 1051-0761. doi: 10.1890/1051-0761(2003)013[0090:CMATEI]2.0.CO;2.

R. S. Cottrell, K. L. Nash, B. S. Halpern, T. A. Remenyi, S. P. Corney, A. Fleming, E. A. Fulton, S. Hornborg, A. Johne, R. A. Watson, and J. L. Blanchard. Food production shocks across land and sea. Nature Sustainability, 2(2):130–137, Feb. 2019. ISSN 2398-9629. doi: 10.1038/s41893-018-0210-1.

F. Courchamp, T. Clutton-Brock, and B. Grenfell. Inverse density dependence and the Allee effect. Trends in Ecology and Evolution, 14(10):405–410, 1999.

L. B. Crowder, S. J. Lyman, W. F. Figueira, and J. Priddy. Source-sink population dynamics and the problem of siting marine reserves. Bulletin of Marine Science, 66(3):23, 2000.

C. C. D’Aloia, R. M. Daigle, I. M. Côté, J. M. Curtis, F. Guichard, and M. J. Fortin. A multiple-species framework for integrating movement processes across life stages into the design of marine protected areas. Biological Conservation, 216(October):93–100, 2017. doi: 10.1016/j.biocon.2017.10.012.

B. Dennis, L. Assas, S. Elaydi, E. Kwessi, G. Livadiotis, and B. Dennis. Allee effects and resilience in stochastic populations. Theoretical Ecology, 9:323–335, 2016. doi: 10.1007/s12080-015-0288-2.

S. Doney, M. Ruckelshaus, J. Emmett Duffy, J. Barry, F. Chan, C. English, H. Galindo, J. Grebmeier, A. Hollowed, N. Knowlton, J. Polovina, N. Rabalais, W. Sydeman, and L. Talley. Climate Change Impacts on Marine Ecosystems. Annual Review of Marine Science, 4(1):11–37, 2012. ISSN 1941-1405\r1941-0611. doi: 10.1146/annurev-marine-041911-111611.

N. S. Fabina, M. L. Baskett, and K. Gross. The differential effects of increasing frequency and magnitude of extreme events on coral populations. Ecological Applications, 25(6): 1534–1545, 2015.

L. Fahrig. Rethinking patch size and isolation effects: The habitat amount hypothesis. Journal of Biogeography, 40(9):1649–1663, 2013. ISSN 1365-2699. doi: 10.1111/jbi.12130.

P. Fiedler. Environmental change in the eastern tropical Pacific Ocean: Review of ENSO and decadal variability. Marine Ecology Progress Series, 244:265–283, 2002. doi: 10.3354/meps244265.

S. D. Gaines, C. White, M. H. Carr, and S. R. Palumbi. Designing marine reserve networks for both conservation and fisheries management. Proceedings of the National Academy of Sciences of the United States of America, 107(43):18286–93, 2010. ISSN 0027-8424. doi: 10.1073/pnas.0906473107.

E. T. Game, E. McDonald-Madden, M. L. Puotinen, and H. P. Possingham. Should we protect the strong or the weak? Risk, resilience, and the selection of marine protected areas. Conservation Biology, 22(6):1619–1629, 2008a. ISSN 1523-1739. doi: 10.1111/j.1523-1739.2008.01037.x.

E. T. Game, M. E. Watts, S. Wooldridge, and H. P. Possingham. Planning for Persistence in Marine Reserves - A Question of Catastrophic Importance. Ecological Applications, 18(3):670–680, 2008b.

L. R. Gerber, L. W. Botsford, A. Hastings, H. P. Possingham, S. D. Gaines, S. R. Palumbi, and S. Andelman. Population Models for Marine Reserve Design : A Retrospective and Prospective Synthesis. Ecological Applications, 13(1):547–564, 2003. ISSN 10510761. doi: 10.1890/1051-0761(2003)013[0047:PMFMRD]2.0.CO;2.

B. S. Halpern and R. R. Warner. Matching marine reserve design to reserve objectives. Proceedings of the Royal Society of London. Series B: Biological Sciences, 270(1527): 1871–1878, Sept. 2003. ISSN 0962-8452, 1471–2954. doi: 10.1098/rspb.2003.2405.

B. S. Halpern, H. M. Regan, H. P. Possingham, and M. A. McCarthy. Accounting for uncertainty in marine reserve design. Ecology Letters, 9(1):2–11, 2006. ISSN 1461-0248. doi: 10.1111/j.1461-0248.2005.00827.x.

A. Hastings and L. W. Botsford. Persistence of spatial populations depends on returning home. Proceedings of the National Academy of Sciences, 103(15):6067–6072, 2006. ISSN 0027-8424. doi: 10.1073/pnas.0506651103.

K. J. Helmstedt, H. P. Possingham, K. E. C. Brennan, J. R. Rhodes, and M. Bode. Cost-efficient fenced reserves for conservation: Single large or two small? Ecological Applications, 24(7):1780–1792, Oct. 2014. ISSN 1051-0761. doi: 10.1890/13-1579.1.

J. A. Hutchings. Renaissance of a caveat: Allee effects in marine fish. ICES Journal of Marine Science, pages 1–6, 2013.

S. Kallimanis, W. E. Kunin, J. M. Halley, and S. P. Sgardelis. Metapopulation Extinction Risk under Spatially Autocorrelated Disturbance. Conservation Biology, 19(2): 534–546, Apr. 2005. ISSN 0888-8892, 1523–1739. doi: 10.1111/j.1523-1739.2005.00418.x.

P. Kinlan and S. D. Gaines. Propagule dispersal in marine and terrestrial environments: A community perspective. Ecology, 84(8):2007–2020, 2003. doi: 10.1890/01-0622.

A. S. Kough, C. B. Paris, D. C. Behringer, and M. J. Butler. Modelling the spread and connectivity of waterborne marine pathogens - the case of PaV1 in the Caribbean. ICES Journal of Marine Science, 72(Supplement 1):i139–i146, 2015. doi: 10.1093/ices.

H. M. Leslie. A Synthesis of Marine Conservation Planning Approaches. Conservation Biology, 19(6):1701–1713, 2005. ISSN 1523-1739. doi: 10.1111/j.1523-1739.2005.00268.x.

S. E. Lester, B. S. Halpern, K. Grorud-Colvert, J. Lubchenco, B. I. Ruttenberg, S. D. Gaines, S. Airamé, and R. R. Warner. Biological effects within no-take marine reserves: A global synthesis. Marine Ecology Progress Series, 384:33–46, May 2009. doi: 10.3354/meps08029.

J. Lubchenco, S. R. Palumbi, S. D. Gaines, S. Andelman, S. E. Applications, S. The, and M. Reserves. Plugging a hole in the ccean: The emerging science of marine reserves. Ecological Applications, 13(1):3–7, 2003.

M. Mangel. On the fraction of habitat allocated to marine reserves. Ecology Letters, 3(1): 15–22, 2000. ISSN 1461-0248. doi: 10.1046/j.1461-0248.2000.00104.x.

M. Mangel and C. Tier. Four facts every conservation biologist should know about persistence. Ecology, 75(3):607–614, 1994.

C. R. McGilliard, A. E. Punt, and R. Hilborn. Spatial structure induced by marine reserves shapes population responses to catastrophes in mathematical models. Ecological Applications, 21(4):1399–1409, 2011.

C. Mellin, M. Aaron Macneil, A. J. Cheal, M. J. Emslie, and M. Julian Caley. Marine protected areas increase resilience among coral reef communities. Ecology Letters, 19 (6):629–637, 2016. ISSN 1461-0248. doi: 10.1111/ele.12598.

K. A. Moloney and S. A. Levin. The Effects of Disturbance Architecture on Landscape-Level Population Dynamics. Ecology, 77(2):375–394, Mar. 1996. ISSN 00129658. doi: 10.2307/2265616.

E. C. Oliver, M. G. Donat, M. T. Burrows, P. J. Moore, D. A. Smale, L. V. Alexander, J. A. Benthuysen, M. Feng, A. Sen Gupta, A. J. Hobday, N. J. Holbrook, S. E. Perkins-Kirkpatrick, H. A. Scannell, S. C. Straub, and T. Wernberg. Longer and more frequent marine heatwaves over the past century. Nature Communications, 9(1):1–12, 2018. doi: 10.1038/s41467-018-03732-9.

R. T. Paine, M. J. Tegner, and E. A. Johnson. Compounded perturbations yield ecological surprises. Ecosystems, 1(July):535–545, 1998. ISSN 1432-9840. doi: 10.1007/s100219900049.

J. W. Pitchford, E. A. Codling, and D. Psarra. Uncertainty and sustainability in fisheries and the benefit of marine protected areas. Ecological Modelling, 207(2-4):286–292, Oct. 2007. ISSN 03043800. doi: 10.1016/j.ecolmodel.2007.05.006.

J. F. Quinn and A. Hastings. Extinction in Subdivided Habitats. Conservation Biology, 1 (3):198–208, 1987.

M. Scheffer and S. R. Carpenter. Catastrophic regime shifts in ecosystems: Linking theory to observation. Trends in Ecology & Evolution, 18(12):648–656, Dec. 2003. doi: 10.1016/j.tree.2003.09.002.

M. Scheffer, S. Carpenter, J. A. Foley, C. Folke, and B. Walker. Catastrophic shifts in ecosystems. Nature, 413(6856):591–6, 2001. ISSN 0028-0836. doi: 10.1038/35098000.

A. F. Shchepetkin and J. C. McWilliams. The regional oceanic modeling system (ROMS): A split-explicit, free-surface, topography-following-coordinate oceanic model. Ocean Modelling, 9(4):347–404, 2005. doi: 10.1016/j.ocemod.2004.08.002.

S. H. Sokolow, P. Foley, J. E. Foley, A. Hastings, and L. L. Richardson. Disease dynamics in marine metapopulations: Modelling infectious diseases on coral reefs. Journal of Applied Ecology, 46(3):621–631, 2009. ISSN 1365-2664. doi: 10.1111/j.1365-2664.2009.01649.x.

L. D. Wagner, J. V. Ross, and H. P. Possingham. Catastrophe management and inter-reserve distance for marine reserve networks. Ecological Modelling, 201(1):82–88, 2007. ISSN 0304-3800. doi: 10.1016/j.ecolmodel.2006.07.030.

J. R. Watson, B. E. Kendall, D. A. Siegel, and S. Mitarai. Changing seascapes, stochastic connectivity, and marine metapopulation dynamics. The American naturalist, 180(1): 99–112, 2012. ISSN 00030147. doi: 10.1086/665992.

E. R. White and A. Hastings. Seasonality in ecology: Progress and prospects in theory. PeerJ Preprints, 7:e27235v2–e27235v2, 2019. doi: 10.7287/peerj.preprints.27235v2.

E. R. White, H. E. Froehlich, J. A. Gephart, R. S. Cottrell, T. A. Branch, and J. K. Baum. Early effects of COVID-19 interventions on US fisheries and seafood. OSF Preprints, 2020. doi: 10.31219/osf.io/9bxnh.

P. D. Williams and A. Hastings. Stochastic dispersal and population persistence in marine organisms. The American naturalist, 182(2):271–82, 2013. ISSN 00030147. doi: 10.1086/671059.

L. J. Wood, L. Fish, J. Laughren, and D. Pauly. Assessing progress towards global marine protection targets: Shortfalls in information and action. Oryx, 42(03):340–351, 2008. ISSN 0030-6053. doi: 10.1017/S003060530800046X.

