## Supplementary Material for "Catastrophes, connectivity, and Allee effects in the design of marine reserve networks"

2020

### Contents

**Appendix S1** ***n*-patch model description** **S2**

**Appendix S2** **Additional figures** **S4**

Data and code for all the figures and tables can be found at (https://github.com/eastonwhite/MPA-disturbances).

### Appendix S1 $n$-patch model description

To extend the two-patch model described in the main manuscript to an $n$-patch scenario, we have to use a slightly different model formulation. We use the same Beverton-Holt structure within each patch for production described in the main manuscript. However, we model spatially-explicit dispersal (i.e. resulting connectivity between patches) and disturbances. We focus on a coastline system (a simple one-dimensional landscape) where $d_{ij}$ is the distance between patches $i$ and $j$ (also see Figure 1d in main text). Thus, the patches are in a contiguous line and where discrete patches next to each other would have a distance of $d_{ij} = 1$ between them.

We use geometric decay for the dispersal kernel, with a dispersal shape parameter $\delta$, where increasing $\delta$ decreases dispersal amount and distance. The probability of dispersal from patch $i$ to $j$ is

$$P(\text{dispersal from patch i to patch j}) = \text{Geometric}(\delta, d_{ij}). \tag{S1}$$

For disturbances, we model the probability of disturbance, $M_i$, in each patch as a binomial process with probability $p_i$:

$$M_i(t) \sim \text{Binomial}(1, p_i). \tag{S2}$$

The spatial extent of the disturbance is a stochastic process giving the disturbance size ($x$), which affects patches near the disturbance. If a disturbance in patch $i$ is larger than the distance between patches $i$ and $j$, $d_{ij}$, then patch $j$ will also be affected by the disturbance:

$$P(\text{disturbance in patch j} \mid \text{disturbance in patch i}) = \begin{cases} 1 & \text{if } d_{ij} < x \\ 0 & \text{if otherwise.} \end{cases} \tag{S3}$$

A disturbance causes density-independent mortality, $\mu$, for the entire patch and all patches with distance $x$.

With this $n$-patch model, we can relax the assumption of a "scorched earth" between patches by setting the fraction of biomass fished in non-reserves to be $F < 1$. This allows us to study the effect of fishing pressure outside reserves on the effectiveness of the marine reserve network.

| Notation | Description | Default value(s) |
| --- | --- | --- |
| $r_i(t)$ | growth factor of patch $i$ at time $t$ described as a normal distribution | |
| $\mu_r$ | mean of growth factor normal distribution | 3 |
| $\sigma_r^2$ | variance of growth factor normal distribution | 0.5 |
| $K_i$ | carrying capacity for patch $i$ | 1 |
| $\omega$ | Allee effect parameter | 1 for no Allee effect or >1 for Allee effect |
| $\delta$ | dispersal kernel shape parameter (larger $\delta$ indicates less dispersal) | 0.7 |
| $p_i$ | probability of disturbance | 0.02 |
| $x$ | size of disturbance (number of patches adjacent to disturbed patch that will also be disturbed) | 1 |
| $\mu$ | density-independent mortality from disturbance | 0.9 |
| $F$ | fraction of biomass fished in non-reserves | for scorched earth assumption of all biomass caught or <1 for moderate levels of fishing |

Table S1: Parameter notation, description, and default values for the n-patch model. As a sensitivity analysis, several parameters are varied in the Figs. 5,6, S4.

### Appendix S2 Additional figures

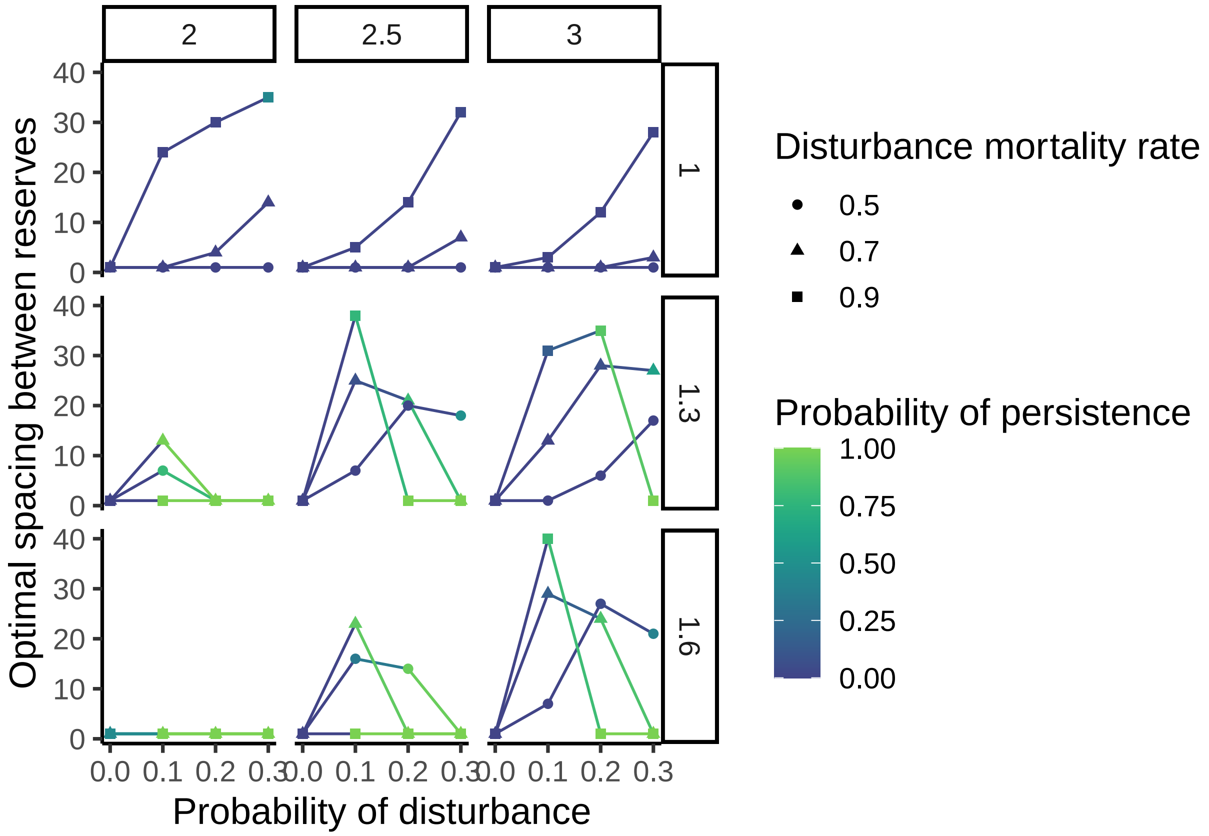


Figure S1: Optimal spacing for varying Allee and r values along with different disturbance parameters.

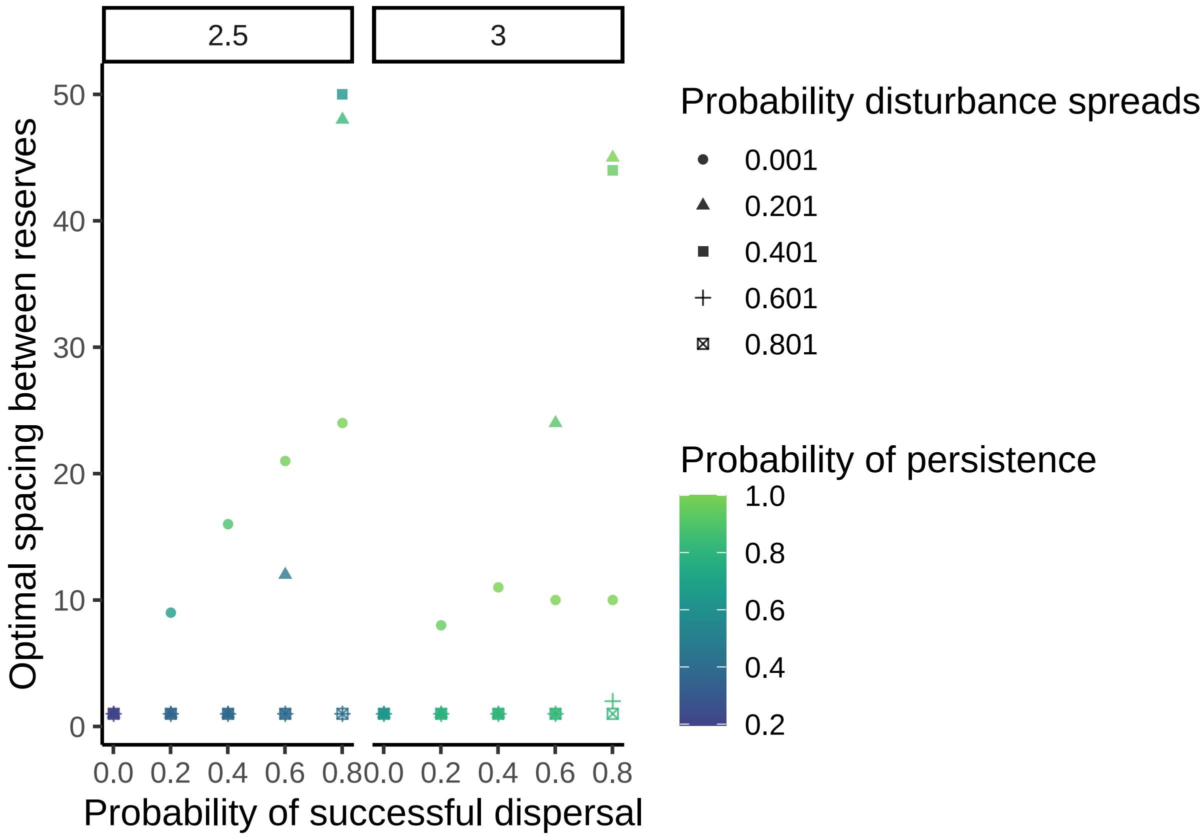


Figure S2: Optimal spacing for different $\gamma$, $\delta$, r, and probability of disturbance.

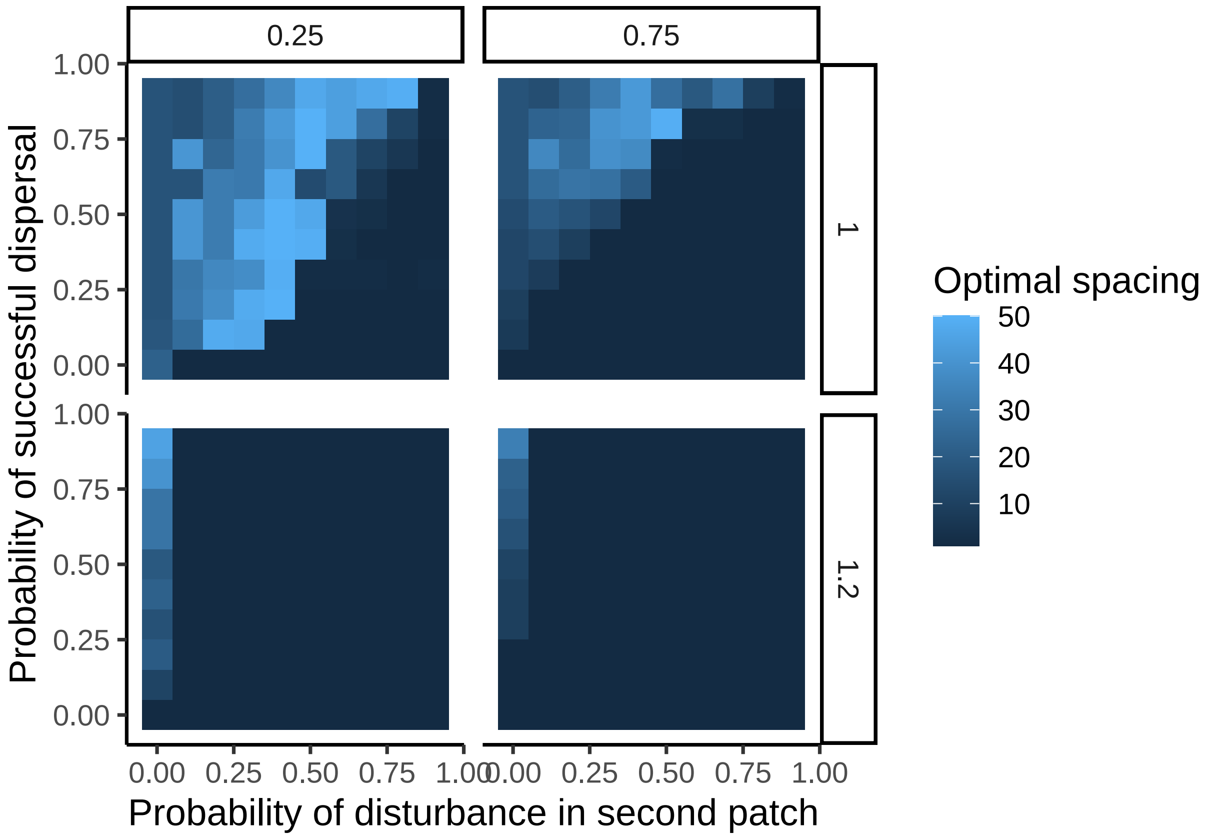


Figure S3: Optimal spacing for all the dispersal parameters and the probability of disturbance.

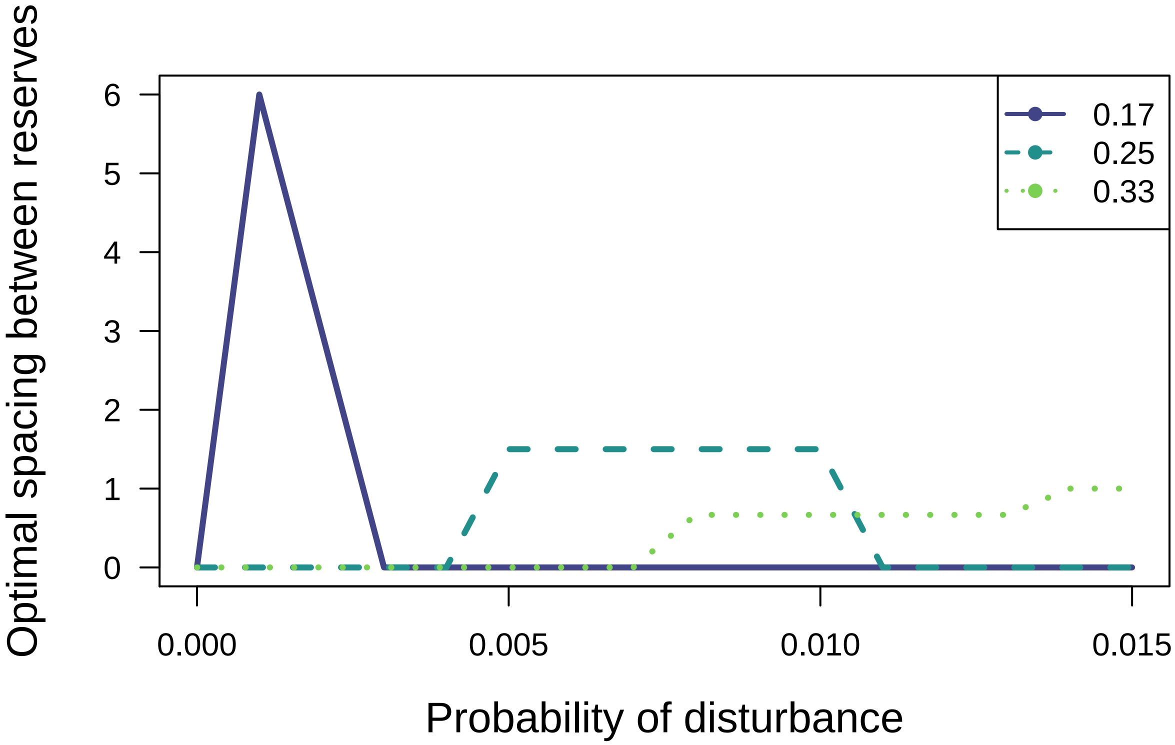


Figure S4: Optimal mean spacing between reserves for different probabilities of disturbance and fraction of coasline in reserves. The specific parameters used here include: $\delta = 0.7$, $\omega = 1.2$, r = 3, and $\mu = 0.9$.
